# Photodynamic therapy in colorectal cancer using photosensitizers functionalized by arene-ruthenium complexes

**DOI:** 10.1101/2022.10.11.511744

**Authors:** Jacquie Massoud, Aline Pinon, Manuel Gallardo-Villagran, Lucie Paulus, Sayed Antoun, Mona Diab-Assaf, Bruno Therrien, Bertrand Liagre

## Abstract

Colorectal cancer (CRC) is one of the most frequently diagnosed cancers worldwide and one of the main causes of cancer deaths in the global population. First-line treatment usually includes surgical procedures followed by combination chemotherapy. Despite the improvements in CRC treatments, the mortality rate remains very high, leading consequently to conventional treatment resistance. Several therapies were enrolled to this content in a way to find the best solutions for this disease. Photodynamic therapy (PDT) using photosensitizers (PS) presents itself as an original innovative therapeutic strategy that strongly limits the undesirable side effects of conventional treatment. However, low physiological solubility and a lack of selectivity of PS towards tumor sites are the principal restrictions of their current clinical use. Indeed, drug delivery systems are currently a key issue in cancer therapy. To overcome the problem of solubility and stability of PS such as porphyrins and their derivatives, metallic assemblies based on arene-ruthenium (Ru) units have begun to attract considerable attention as PS delivery systems. The purpose of this study was to demonstrate firstly the interest in the vectorization of tetrapyridylporphin (TPyP-arene-Ru) and Zn-tetrapyridylporphin arene-ruthenium (Zn-TPyP-arene-Ru) metallacages to increase their solubility in biological media and then, consequently their anticancer efficacy in PDT. Secondly, to elucidate the anticancer mechanism as well as identify the cell death process mediated by these new vectorized PS. The results showed that the two PS-arene-Ru complexes have a strong photocytotoxic effect after photoactivation on human HCT116 and HT-29 colorectal cancer cell lines. TPyP-arene-Ru-PDT induced outstanding cytotoxicity when compared to the Zn-TPyP-arene-Ru analogue. The two complexes show no significant effect on proliferation in the dark. In addition, results demonstrated that these two PS-arene-Ru complexes-PDT induce an apoptotic process through the appearance of a sub-G1 peak, phosphatidylserines externalization, poly-ADP ribose polymerase (PARP) cleavage and DNA fragmentation. Thus, our data contribute to highlighting that the incorporation of porphyrins in Ru-based assemblies could be an efficient vectorized system to treat CRC by PDT.

## 1. Introduction

Cancer is a group of diseases that refer to abnormal cell division leading to uncontrolled cell growth and proliferation. When this type of growth occurs in the colon or rectum, this disease is defined as colorectal cancer (CRC) [1]. In 2022, the World Health Organization (WHO) and the International Agency for Research on Cancer (IARC) estimate that in 2020, CRC is the third most common cancer worldwide with approximately 1.9 million cases annually and the second leading of death for oncological reasons globally with 900 000 deaths [2,3]. CRC mainly originate from a benign tumor or adenomatous polyp that evolves into a dangerous malignant tumor [4]. The passage from a benign tumor to a malignant one is characterized by the capacity of the cells to infiltrate the different histological layers of the organ [5]. The most dangerous stage is the metastatic stage when the cancer cells have acquired the ability to detach from the initial tumor and invade other organs, through the blood or lymph, and create a secondary tumor [6]. Today, CRC is at a crossroads, strategies used to treat it involve many conventional and advanced scientific methods. These therapies incorporate surgery/polypectomy, radiation therapy, chemotherapy, combination therapies, and targeted therapy [7–9]. Unfortunately, these methods are not sufficient for the complete treatment of CRC. Researchers have tried to provide advanced alternative approaches to deal with conventional methods’ drawbacks and combat CRC resistance to conventional treatments [10]. Over the past decade, significant progress in CRC treatment has been achieved through the development of novel drugs and treatment protocols. However, the increasing resistance of tumor cells toward these novel drugs and persistent side effects due to their toxicity on healthy tissues make it imperative to find other methods for CRC therapy.

Photodynamic therapy (PDT) has attracted widespread attention in recent years as a non-invasive and highly selective approach for cancer treatment precisely for CRC. The molecular mechanism of PDT involves the photoactivation of a photosensitizer (PS) by an appropriate wavelength of light in presence of oxygen molecules in the tumor environment [11–13]. In effect, PDT exploits the potency of visible light to deliver cytotoxic agents in a spatially and temporally controlled manner to directly damage the targeted tumor cells and tissues [14,15]. PDT is mainly targeted toward the generation of reactive oxygen species (ROS) which are involved in cell toxicity [12,16,17]. To occur photodynamic photochemical reactions, the PS must absorb at least one photon to be promoted to a sufficiently long-lived excited state and then induce photodynamic reactions in an oxygenated environment [18]. Under the effect of light irradiation, the PS is activated and goes from the ground state to an excited one [19,20]. At this stage, the PS is very unstable and loses its excess energy either directly or via the excited triplet state [21]. The excited triplet will slowly return to the ground state by photochemical reactions of type I or II. Both reactions may take place simultaneously being their kinetics strongly favored by the oxygen, substrate concentration, and type of the PS. In type I reaction, the free radicals may further react with oxygen to produce ROS [22,23]. Superoxide anion initially produced via type I pathway by monovalent reduction does not cause oxidative damage but reacts with itself to generate oxygen and hydrogen peroxide. However, in type II reaction, the excited PS transfers its energy directly to molecular oxygen to form singlet oxygen. These highly cytotoxic ROS can oxidize a variety of biomolecules, inducing an acute stress response and triggering a series of redox signaling pathways, frequently leading to cell death [24–26].

Currently, the most widely PS used in PDT are the tetrapyrrole derivate such as porphyrins, chlorines, bacteriochlorines, and phthalocyanines. Nevertheless, the main inconveniences of these PS are their low water solubility, which limits intravenous administration, their poor photophysical properties due to PS aggregation, and their low tumor selectivity, limiting consequently their use in clinical trials [27]. Constantly, the most important aim of the research today is to find the best PS, overcome their solubility problems, and improve their effectiveness in PDT.

The reactivity and interactions theory between metal complexes and biomolecules such as DNA and proteins has made them the center of interest for therapeutic purposes. Unfortunately, this type of drug has shown significant side effects, which has led to the search for new, less toxic anticancer agents [28]. Ruthenium (Ru) complexes with polypyridyl ligands have received much attention due to their interesting properties [29,30]. These properties lead to their potential use in various fields such as PS for photochemical conversion and precisely as photoactive DNA cleavage agents for therapeutic purposes. Porphyrins and their derivatives are one of the outstanding PS conjugated to Ru [31,32]. Indeed, the conjugation of porphyrins to peripheral metal moieties is an interesting strategy for the development of compounds that may combine the cytotoxicity of the metal moiety with the phototoxicity of the porphyrin chromophore for additive antitumor effects [33,34]. To circumvent the problem of solubility and stability of Ru-based PS, metal assemblies have begun to attract considerable interest [35]. These molecular assemblies have been used to generate an isolated environment for PS as encapsulation systems and protect it as guest, sensitive or unstable molecules. These assemblies are known as “host-guest” systems [36]. Our collaborators Gallardo-Villagrán et al. reported by several studies that the poor solubility of some PS could be solved by using arene-Ru-based metallacages as carriers in two different ways [37]. The first type refers to a prismatic metallacages, which can improve the solubility of these organometallic complexes in biological media, likewise, they can entertain PS as guest in their internal cavities, transporting and releasing them into the target cells [38]. The second type of this vectorization system is cubic metallacages. The specificity of these metallacages relies on the integration of the PS into the initial structure of the metallacages, which maintains an excellent physiological solubility and keeps the PS available to be irradiated at any time [39].

In the ongoing study, we focused on the evaluation of the anticancer efficacy of the cubic arene-Ru metallacages carrying on their own structure two units of PS based on tetrapyridylporphin (TPyP) as an interesting vectorization system. In addition, to demonstrate a possible metal influence on the photoactivity of the PS, we examined the presence of diamagnetic metal (Zn^2+^) in the center of the tetrapyrrole ring in the TPyP panels (Zn-TPyP). In such a way, we verified whether the photoactivity of the PS has been modified or not when it is part of a metal assembly. Firstly, we investigate the anticancer effect of the functionalized TPyP and Zn-TPyP with arene-Ru complexes on human HCT116 and HT-29 colorectal cancer cell lines. Then, for purpose of understanding the cell death process involved in this therapy, we examined the cell cycle distribution, phosphatidylserines externalization, as well as poly-ADP ribose polymerase (PARP) cleavage, and DNA fragmentation. Subsequently, consistent with other PDT studies, our results demonstrated that TPyP and Zn-TPyP-arene-Ru complexes (Figures 1 and 2) achieve their anticancer effects through the apoptotic process.

**Figure 1:**
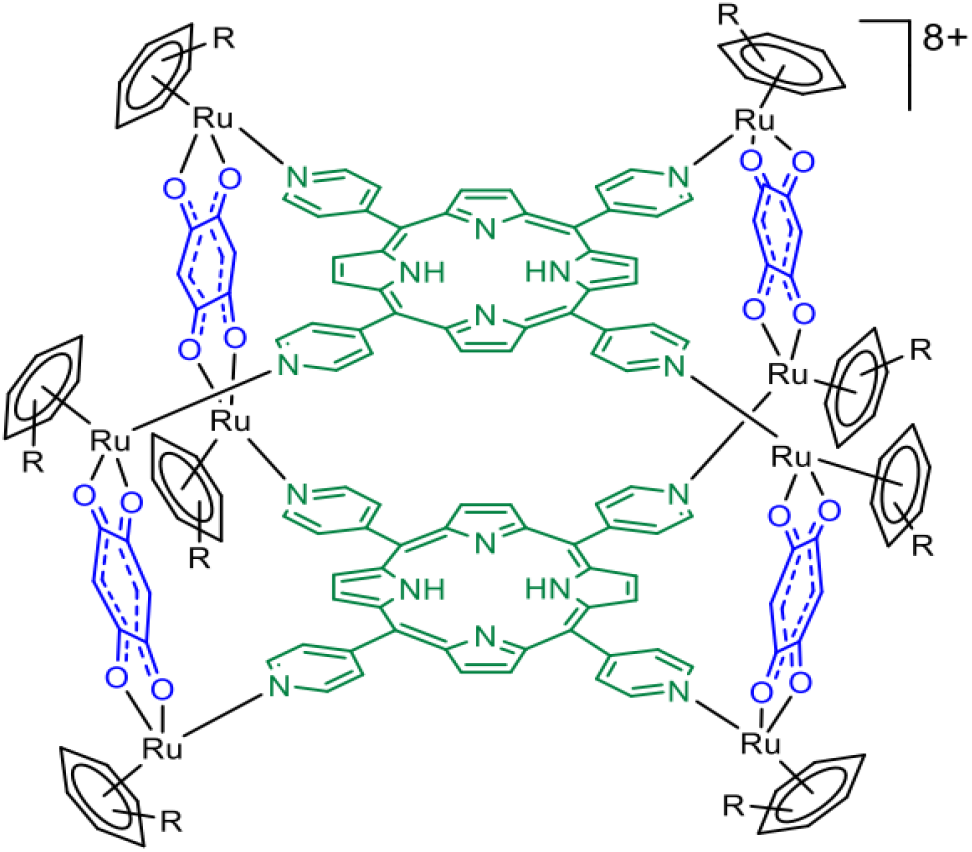
Ruthenium metallacages containing TPyP.

**Figure 2:**
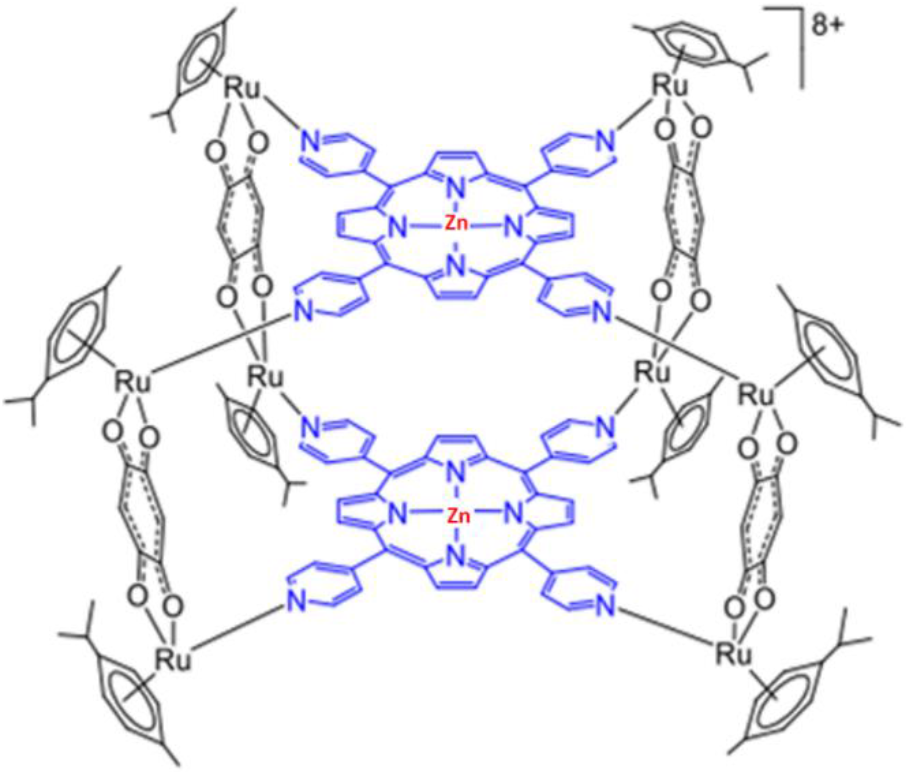
Ruthenium metallacages containing Zn-TPyP.

## 2. Material and Methods

### 2.1. Materials

DMEM medium, DMEM red-phenol-free medium, RPMI 1640 medium, RPMI 1640 red-phenol-free medium, fetal bovine serum (FBS), L-glutamine, and penicillin-streptomycin were purchased from Gibco BRL (Cergy-Pontoise, France). 3-(4,5-dimethylthiazol-2-yl)-2,5-diphenyltetrazolium bromide (MTT), cell death detection enzyme-linked immunosorbent assay ^PLUS^ (ELISA) and human anti-β-actin antibody were obtained from Sigma-Aldrich (Saint-Quentin-Fallavier, France). Poly-ADP-ribose polymerase (PARP) antibody and goat anti-rabbit IgG secondary antibody conjugated to horseradish peroxidase (HRP) were acquired from Cell Signaling Technology-Ozyme (Saint-Quentin-en-Yvelines, France). Rabbit anti-mouse IgG-IgM H&L HRP secondary antibody, Annexin V-FITC, and propidium iodide (PI) were obtained from Invitrogen-Thermo Fisher Scientific (Villebon-Sur-Yvette, France). Immobilon Western Chemiluminescent HRP Substrate was acquired from Merck (Lyon, France).

### 2.2. Cell culture and treatment

Human CRC cell lines HT-29 and HCT116 were purchased from the American Type Culture Collection (ATCC-LGC Standards, Mosheim, France). Cells were grown in DMEM medium for HT-29 cells and RPMI 1640 medium for HCT116 cells. Culture mediums were supplemented with 10% FBS, 1% L-glutamine and 100 U/mL penicillin, and 100 μg/mL streptomycin. Cultures were maintained in a humidified atmosphere containing 5% CO_2_ at 37°C. For all experiments, cells were seeded at 2.1×10^4^, 1.2×10^4^ cells/cm^2^ for HT-29 and HCT116 cells respectively and the culture medium was replaced by a red phenol-free appropriate culture medium before PDT.

TPyP and Zn-TPyP-arene-Ru complexes were prepared as previously described [40–42]. Ru (III) chloride hydrate, 1-isopropyl-4-methylcyclohexane-1,4-diene, 5,10,15,20-tetra(4-pyridyl)-21*H*,23*H*-porphine and zinc 5,10,15,20-tetra(4-pyridyl)-21*H*,23*H*-porphine were acquired from Sigma-Aldrich. Stock solutions of PS-arene-Ru complexes were dissolved at 1 mM in DMSO, then were diluted in culture medium to obtain the appropriate final concentrations just before use. The concentration of DMSO in culture medium was lower than 0.1% in all cases.

### 2.3. Phototoxicity of TPyP and Zn-TPyP-arene-Ru complexes

Phototoxicity was determined using an MTT assay. Briefly, cells were seeded in 96-well culture plates and grown for 36h in an appropriate culture medium before exposure or not to TPyP or Zn-TPyP-arene-Ru complexes. After 24h incubation, cells were irradiated or not with a 630-660 nm CURElight lamp at 75 J/cm^2^ (PhotoCure ASA, Oslo, Norway). MTT assays were performed 24 and 48h post-irradiation and cell viability was expressed as a percentage of each treatment condition by normalizing to untreated cells.

### 2.4. Cell cycle analysis

The cell cycle distributions in colorectal HT-29 and HCT116 cell lines were analyzed by flow cytometry using propidium iodide (PI) staining. For each cell line, cells were treated or not with the determined IC_50_ values of TPyP and Zn-TPyP-arene-Ru for 24 and 48h, then harvested with trypsin. For flow cytometry analysis, 1.5×10^6^ cells of each condition were collected, washed with PBS, and fixed by adding 1 mL of chilled 70% ethanol in PBS and stored at −20°C. Following fixation, cells were pelleted, washed in cold PBS, resuspended in 500 mL of cold PBS containing 30 μL of RNase A (10 mg/mL), and incubated for 30 min at room temperature. After staining with 25 μL of PI, the percentage of cells in each stage of the cell cycle was determined using the FACS system (BD Biosciences, San Jose, CA). All the experiments were performed on three samples.

### 2.5. Apoptosis by TPyP and Zn-TPyP-arene-Ru-PDT

#### 2.5.1. Annexin V-FITC/PI dual staining assay

The Annexin V-FITC/PI dual staining assay was used to determine the percentage of apoptotic cells. For each cell line, cells were treated or not with the determined IC_50_ values of TPyP and Zn-TPyP-arene-Ru for 24 and 48h and then harvested with trypsin. 2.5×10^5^ cells of each condition were collected, washed in PBS, centrifuged, and resuspended in 100 μL binding buffer (1X) containing 5 μL of Annexin V-FITC and 1 μL of PI (0.1 mg/mL) at room temperature in the dark. After 15 min incubation, cells were analyzed for the percentage undergoing apoptosis using the FACS system (BD Biosciences). All the experiments were performed on three samples.

#### 2.5.2. Protein extraction and western blot analysis

For each cell line, cells were treated or not with the determined IC_50_ values of TPyP and Zn-TPyP-arene-Ru for 24 and 48h and then harvested with trypsin. For total protein extraction, collected samples of each condition were washed in PBS. Then, the total cell pool was centrifuged at 200 g for 5 min at 4°C and homogenized in RIPA lysis buffer (50 mM HEPES, pH 7.5, 150 mM NaCl, 1% sodium deoxycholate, 1% NP-40, 0.1% SDS, 20 mg/mL of aprotinin) containing protease inhibitors according to the manufacturer’s instructions as previously described [43]. The protein level was determined using the Bradford method. Proteins (60 μg) were separated on 12.5% SDS-PAGE gels and transferred to PVDF membranes (Amersham Pharmacia Biotech, Saclay, France). Membranes were probed with respective human antibodies against PARP and β-actin used as a loading control, according to the manufacturer’s instructions. After incubation with appropriate secondary antibodies, blots were developed using the “Immobilon Western” substrate following the manufacturer’s protocol and G: BOX system (Syngene, Ozyme).

#### 2.5.3. Apoptosis quantification: DNA fragmentation

For each cell line, cells were treated or not with the determined IC_50_ values of TPyP and Zn-TPyP-arene-Ru for 24 and 48h and then harvested with trypsin. Histone release from the nucleus during apoptosis was analyzed using the Cell Death Detection ELISA^PLUS^ as previously described [44]. 2×10^5^ cells of each condition were obtained and DNA fragmentation was measured according to the manufacturer’s protocol.

#### 2.5.4. Statistical Analysis

All quantitative results are expressed as the mean ± standard error of the mean (SEM) of separate experiments. Statistical significance was evaluated by the two-tailed unpaired Student’s t-test and expressed as: * p < 0.05; ** p < 0.01 and *** p < 0.001.

## 3. Results

### 3.1. Phototoxic effect of TPyP and Zn-TPyP-arene-Ru

To investigate the *in vitro* phototoxicity of TPyP and Zn-TPyP-arene-Ru, we treated or not two human CRC cell lines: HCT116 and HT-29 with TPyP or Zn-TPyP-arene-Ru complexes. Then, the cells were exposed or not to PDT with red irradiation and phototoxic effects were determined 24 and 48h post-irradiation using the MTT assay. Results showed that PS-arene-Ru complexes had no toxic effect on HCT116 and HT-29 cell lines when the cells were kept in the dark. Forwards PDT activation, results revealed that both TPyP and Zn-TPyP-arene complexes leads to a drastic decrease in cell viability in a dose-dependent manner (Figures 3 and 4). However, TPyP-arene-Ru complex was more effective than the Zn-TPyP analogue on both cell lines.

**Figure 3:**
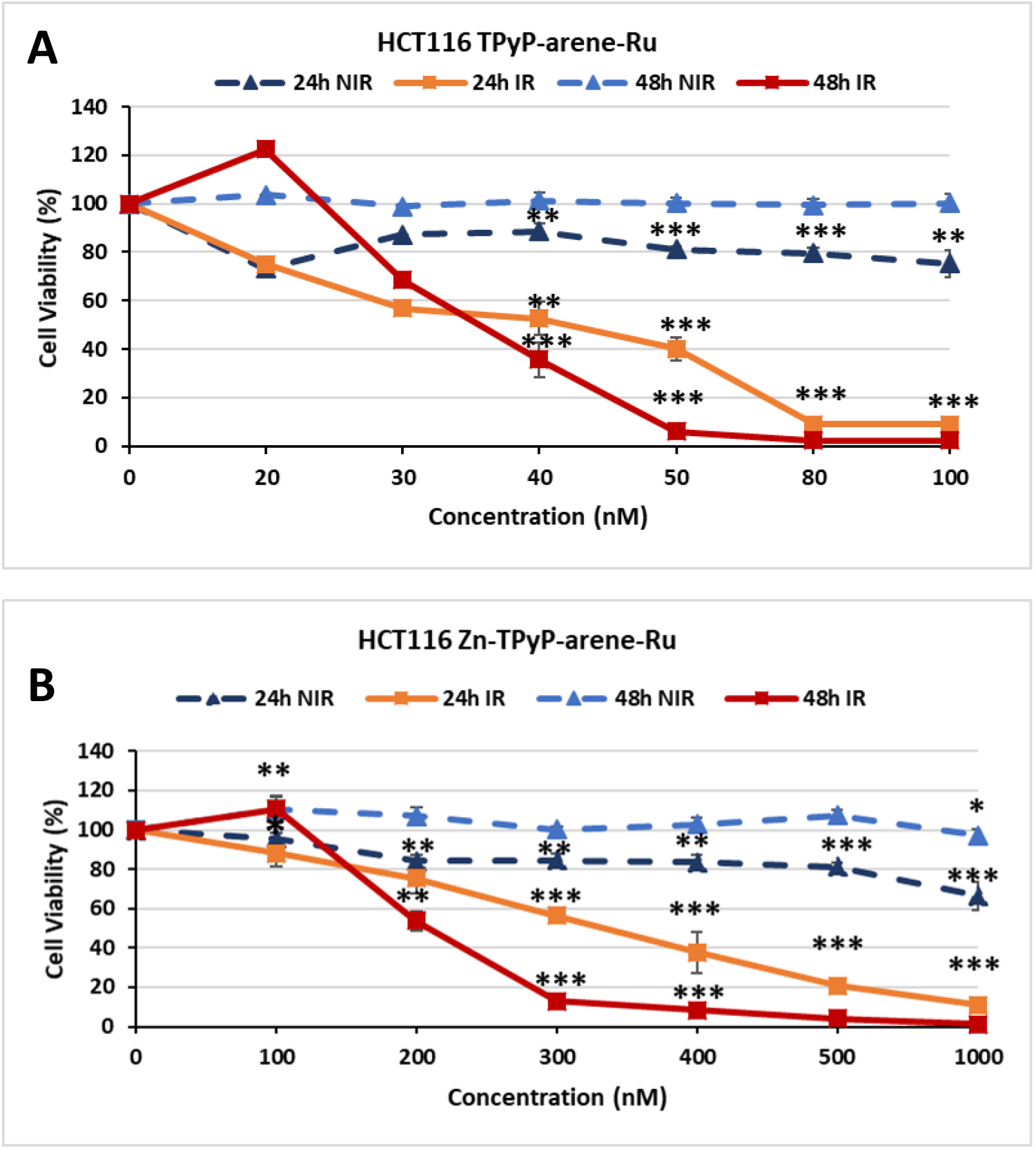
Photocytotoxic effect of TPyP-arene-Ru (A) and Zn-TPyP-arene-Ru (B) on HCT116 cells. Cells were seeded in 96-well culture plates and grown for 36h in an appropriate culture medium before exposure or not to TPyP or Zn-TPyP-arene-Ru complexes. After 24h incubation, cells were irradiated or not with a 630-660 nm CURElight lamp at 75 J/cm^2^ (PhotoCure ASA, Oslo, Norway). MTT assays were performed 24 and 48h post-irradiation and cell viability was expressed as a percentage of each treatment condition by normalizing to untreated cells. Data are shown as mean ± SEM (n = 3). * *p* < 0.05, ** *p* < 0.01 and \*\*\**p* < 0.001.

**Figure 4:**
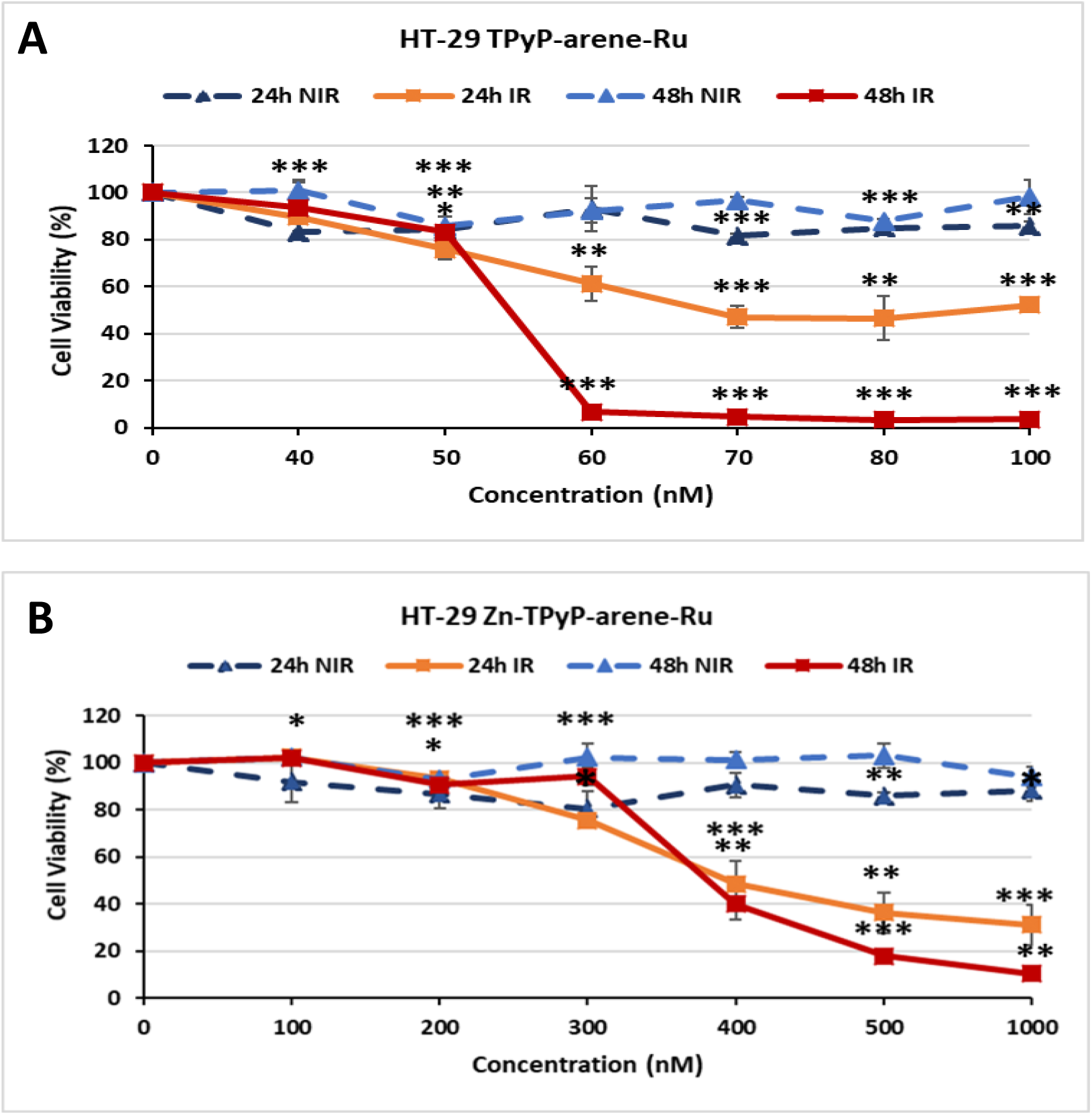
Photocytotoxic effect of TPyP-arene-Ru (A) and Zn-TPyP-arene-Ru (B) on HT-29 cells. Cells were seeded in 96-well culture plates and grown for 36h in an appropriate culture medium before exposure or not to TPyP or Zn-TPyP-arene-Ru complexes. After 24h incubation, cells were irradiated or not with a 630-660 nm CURElight lamp at 75 J/cm^2^ (PhotoCure ASA, Oslo, Norway). MTT assays were performed 24 and 48h post-irradiation and cell viability was expressed as a percentage of each treatment condition by normalizing to untreated cells. Data are shown as mean ± SEM (n = 3). * *p* < 0.05, ** *p* < 0.01 and \*\*\**p* < 0.001.

IC_50_ values were determined to compare the impact of adding a diamagnetic metal (Zn^2+^) to the center of the tetrapyrrole ring in TPyP panels after activation with PDT. We observed that TPyP-arene-Ru was much more effective than Zn-TPyP-arene-Ru in the HCT116 cell line with 8-fold more cytotoxicity at 24h (Figure 3A and 3B). IC_50_ values were in range of 41.89 nM for TPyP-arene-Ru and 331.23 nM for Zn-TPyP-arene-Ru. Our compounds revealed more cytotoxicity at 48h where IC_50_ values decreased to 35.24 nM for TPyP-arene-Ru and 207.45 nM for Zn-TPyP-arene-Ru (Figure 3, Table 1).

**Table 1:** IC_50_ values determined with MTT assays on HCT116 cells.

| PS | IC<sub>50</sub>/PDT<br>24h | IC<sub>50</sub>/PDT<br>48h | IC<sub>50</sub>/Obscurity |
| --- | --- | --- | --- |
| TPyP-arene-Ru | 41.89 | 35.24 | > 100 |
| Zn-TPyP-arene-Ru | 331.23 | 207.45 | > 1000 |

Similar results were noted for the HT-29 cell line that showed more resistant than HCT116 as their respective IC_50_ values were 67.82 nM for TPyP-arene-Ru and 393.89 nM for Zn-TPyP-arene-Ru at 24h (Figure 4A and 4B). These values decreased to be respectively 54.10 nM for TPyP-arene-Ru and 379.29 nM for Zn-TPyP-arene-Ru at 48h (Figure 4, Table 2).

**Table 2:** IC_50_ values (nM) determined with MTT assays on HT-29 cells.

| PS | IC<sub>50</sub>/PDT<br>24h | IC<sub>50</sub>/PDT<br>48h | IC<sub>50</sub>/Obscurity |
| --- | --- | --- | --- |
| TPyP-arene-Ru | 67.82 | 54.10 | > 100 |
| Zn-TPyP-arene-Ru | 393.89 | 379.29 | > 1000 |

The compounds were considered at the determined IC_50_ values for the following experiments.

### 3.2. TPyP and Zn-TPyP-arene-Ru effect on cell cycle

PDT can induce irreversible photodamage leading to cell death. To define the cell death process triggered by our compounds and to confirm that the viability reduction in the two CRC cell lines studied was due to the photoactivation of TPyP and Zn-TPyP-arene-Ru complexes by PDT, we decided to verify the changes in the cell cycle induced by this photoactivation.

HCT116 and HT-29 cells were treated or not with the determined IC_50_ values of TPyP or Zn-TPyP-arene-Ru complexes and then subjected to flow cytometry analysis using PI staining after PDT. Results showed that on HCT116 cells TPyP-arene-Ru induced a strong increase in the number of apoptotic cells represented by the sub-G1 peak mainly at 48h within 33.83% vs 0.65% for control. In contrast, Zn-TPyP-arene-Ru was shown to have less effect on sub-G1 cells with 5.29% vs 0.65% for control at 48h (Figure 5). Similar results were observed for HT-29 cells where TPyP-arene-Ru produced an increase in the number of apoptotic cells as appeared by a sub-G1 peak with 9.04% vs 1.22% for the control at 24h. Likewise, we observed a drastic increase at 48h with a 28.30% sub-G1 peak vs 1.22% for the control (Figure 6). It is important to note that our two complexes have no influence on the cell cycle in the dark in both cell lines tested.

**Figure 5:**
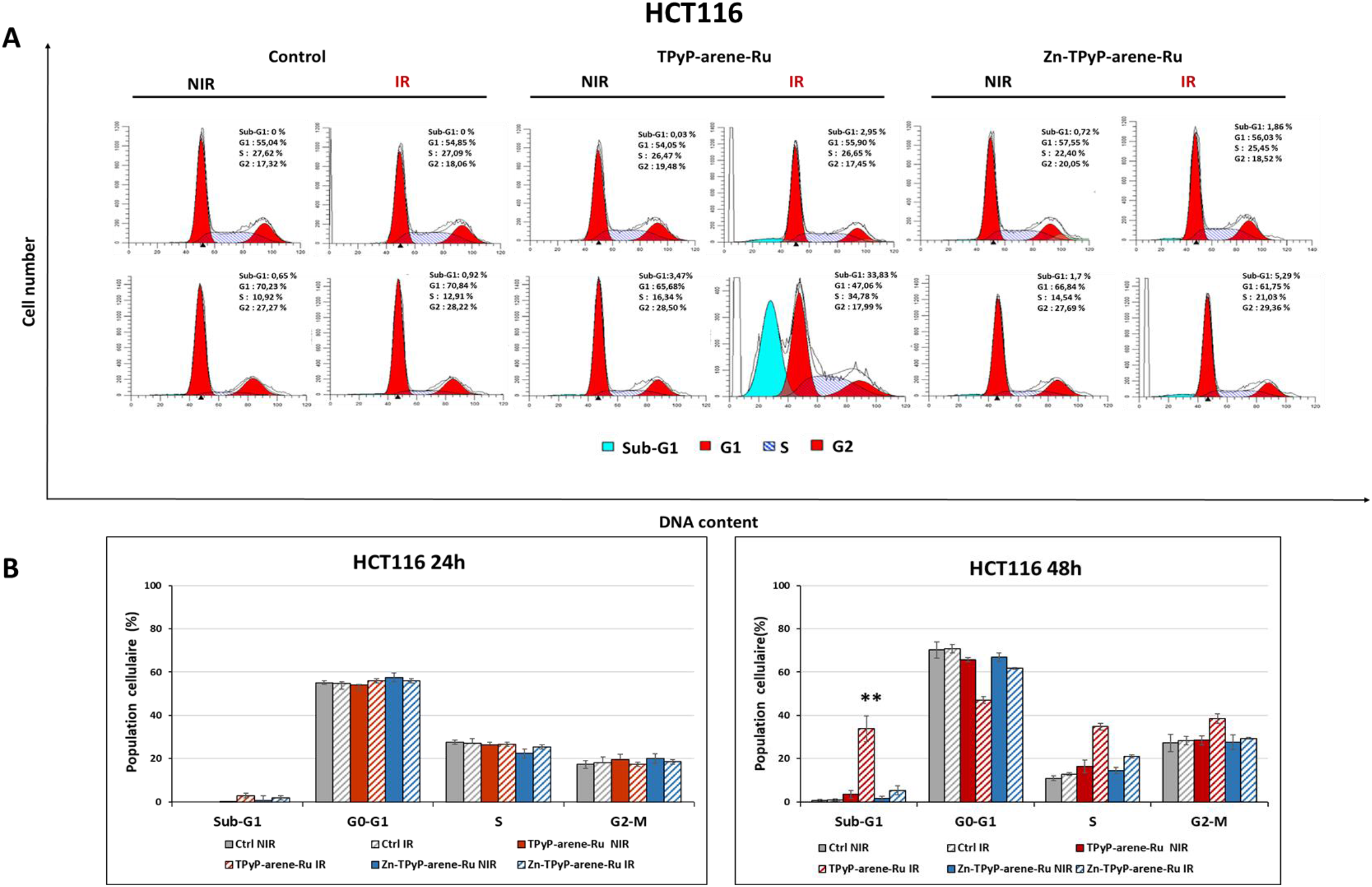
Effects of photoactivation of PS-arene-Ru on the cell cycle distribution in HCT116 cells. Cells were grown for 36h in an appropriate culture medium before exposure or not to TPyP or Zn-TPyP-arene-Ru complexes. After 24h incubation, cells were irradiated or not with a 630660 nm CURElight lamp at 75 J/cm^2^ (PhotoCure ASA, Oslo, Norway). Then subjected to flow cytometry analysis using PI staining after PDT. Images of cell cycle analysis (A) were representative of three separate experiments. Results of flow cytometry analysis are represented by histograms (B) that display the percentage of cells in each cell cycle phase. Data are shown as mean ± SEM (n = 3). *** *p* < 0.001

**Figure 6:**
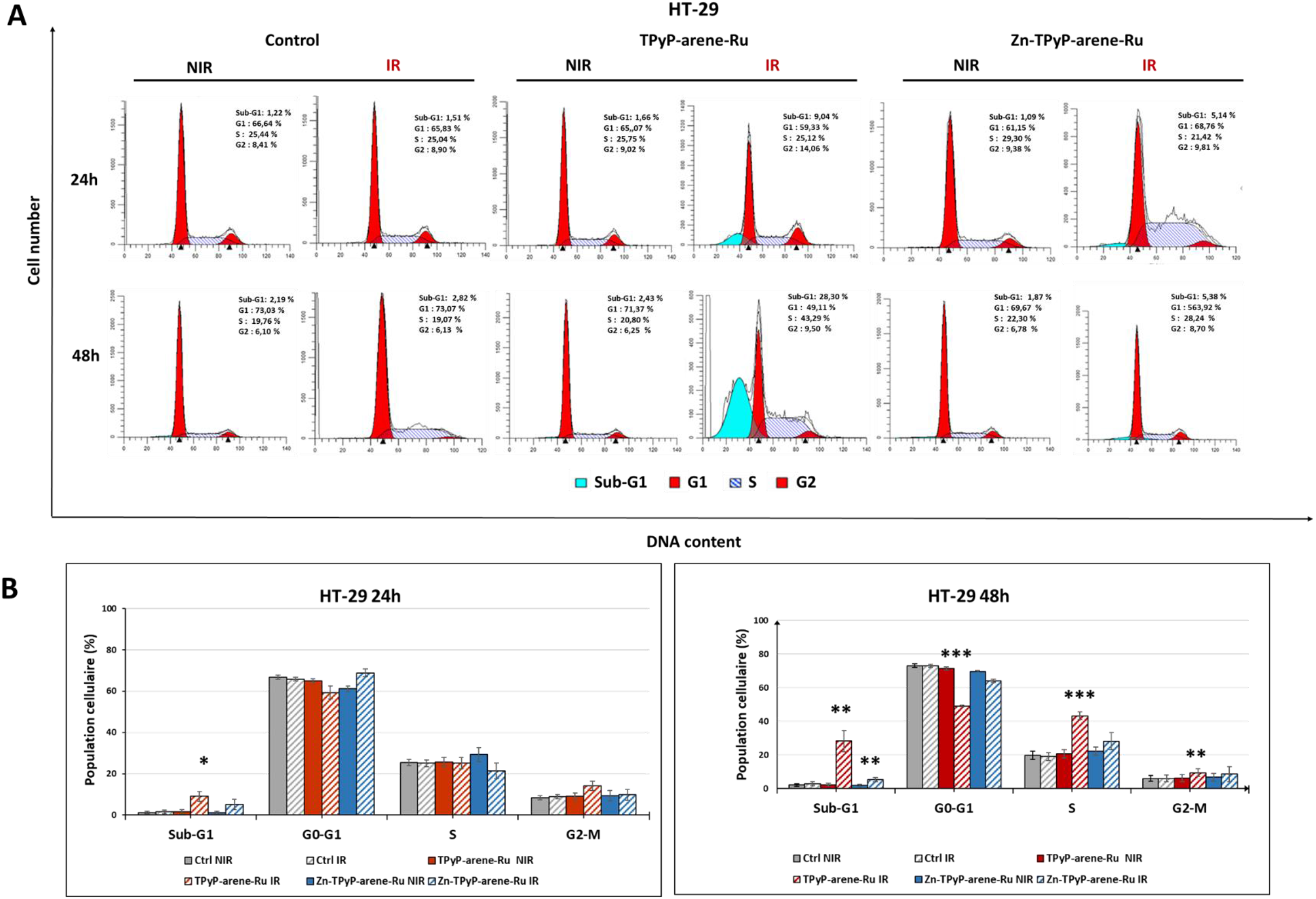
Effects of photoactivation of PS-arene-Ru on the cell cycle distribution in HT-29 cells. Cells were grown for 36h in an appropriate culture medium before exposure or not to TPyP or Zn-TPyP-arene-Ru complexes. After 24h incubation, cells were irradiated or not with a 630-660 nm CURElight lamp at 75 J/cm^2^ (PhotoCure ASA, Oslo, Norway). Then subjected to flow cytometry analysis using Propidium Iodide (PI) staining after PDT. Images of cell cycle analysis (A) shown were representative of three separate experiments. Results of flow cytometry analysis are represented by histograms (B) that display the percentage of cells in each cell cycle phase. Data are shown as mean ± SEM (n = 3). \**p* < 0.05, \*\**p* < 0.01 and *** *p* < 0.001.

### 3.3. TPyP and Zn-TPyP-arene-Ru induce apoptosis

Cell cycle analysis of HCT116 and HT-29 cells treated with either TPyP or Zn-TPyP-arene-Ru revealed the appearance of a sub-G1 population referring to cells in apoptosis. Therefore, we have studied the mechanism of apoptosis induced by TPyP and Zn-TPyP-arene-Ru complexes on HCT116 and HT-29 cells 24 and 48h post-PDT. The apoptotic process was first investigated using annexin V-FITC/PI dual staining assay. During the early stages of apoptosis, phosphatidylserines are known for their translocation from the inner to the outer plasma membrane of the cell, thus, phosphatidylserines externalization allows it to bind with annexin V. Therefore, the percentages of apoptotic cells at early and later stages were determined by dual staining with Annexin V-FITC and PI by flow cytometry. Results showed that in HCT116 cells, control, light control, TPyP and Zn-TPyP-arene-Ru treated cells were mostly viable, whereas the cumulative rate of early and late apoptosis was 11.31%, 11.29%, 9.79%, and 10.23% respectively at 24h. This rate has increased dramatically with the conjugation of irradiation to be 49.25% for TPyP-arene-Ru-PDT more effective than Zn-TPyP-arene-Ru-PDT with 21.58%. Same as for 48h after PDT, TPyP-arene-Ru and Zn-TPyP-arene-Ru caused 61.46% and 42.38% of apoptosis respectively compared to 12.96% for control, 15.24% for light control, 16.22% for TPyP-arene-Ru and 22.4% for Zn-TPyP-arene-Ru complexes without photoactivation (Figure 7).

**Figure 7:**
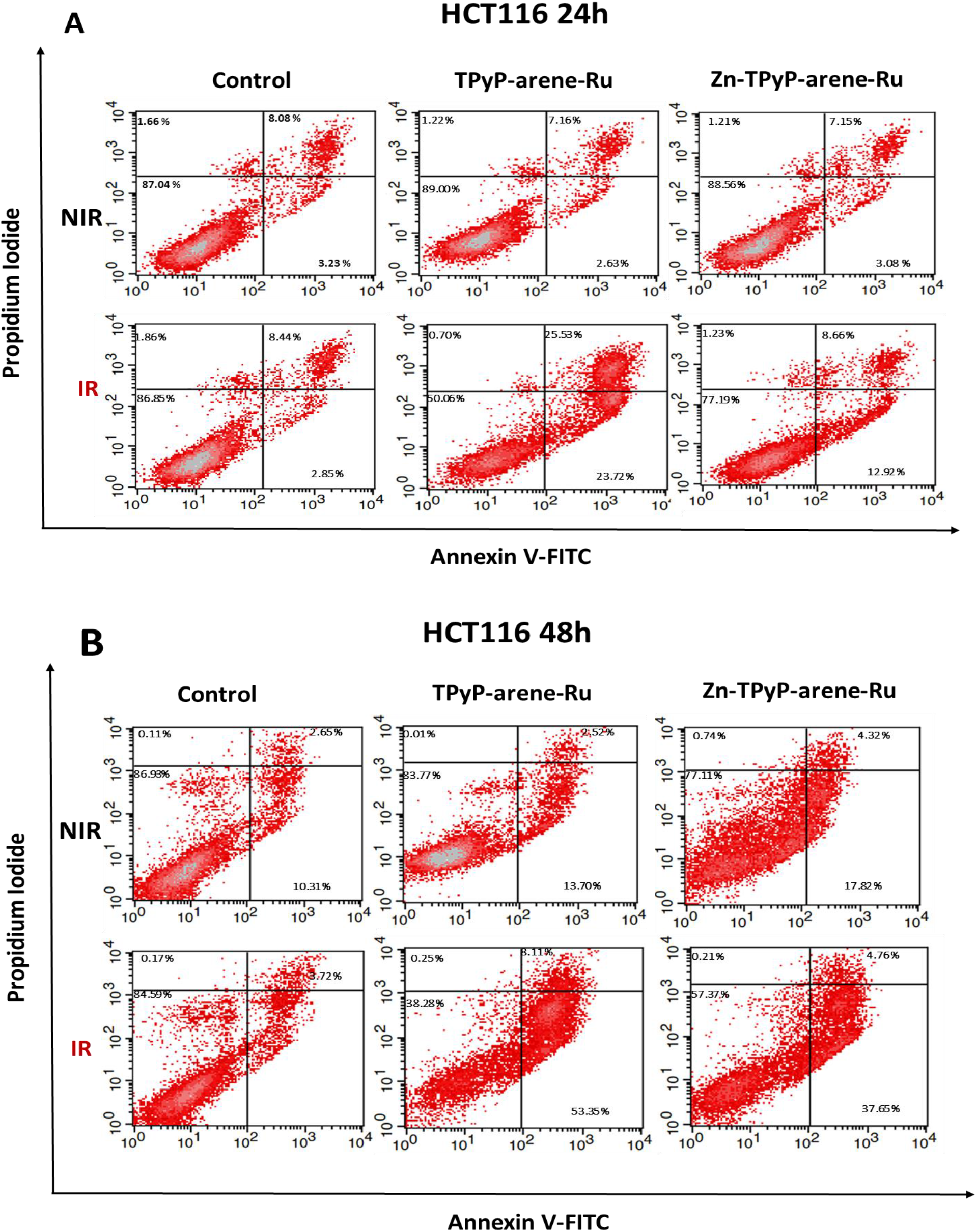
Effects of photoactivation of PS-arene-Ru on HCT116 cell line apoptosis. Cells were grown for 36h in an appropriate culture medium before exposure or not to TPyP or Zn-TPyP-arene-Ru complexes. After 24h incubation, cells were irradiated or not with a 630-660 nm CURElight lamp at 75 J/cm^2^ (PhotoCure ASA, Oslo, Norway). HCT116 cells were also stained, 24h post-PDT (A) and 48h-post-PDT (B) with Annexin V-FITC and PI, and apoptosis was analyzed by flow cytometry. The upper right quadrant represents the percentage of late apoptosis, and the lower right quadrant represents early apoptosis. Data are shown as mean ± SEM (n = 3).

HT-29 cells showed to be less sensitive than HCT116 cells. At 24h, control, light control, TPyP and Zn-TPyP-arene-Ru treated cells revealed as expected a high viability percentage with apoptosis total rate of 6.7%, 8.02%, 8.38%, and 7.29%. However, when TPyP and Zn-TPyP-arene-Ru complexes were photoactivated, they increase the cumulative rate of early and late apoptosis to 35.45% and 22.8% respectively. In addition, 48h post-irradiation, TPyP and Zn-TPyP-arene-Ru complexes strongly increased the apoptosis rate to 58.31% and 36.82% respectively. Control, control light, TPyP and Zn-TPyP-arene-Ru complexes showed 13.44%, 12.92%, 10.67% and 13.07% as total apoptosis rate (Figure 8).

**Figure 8:**
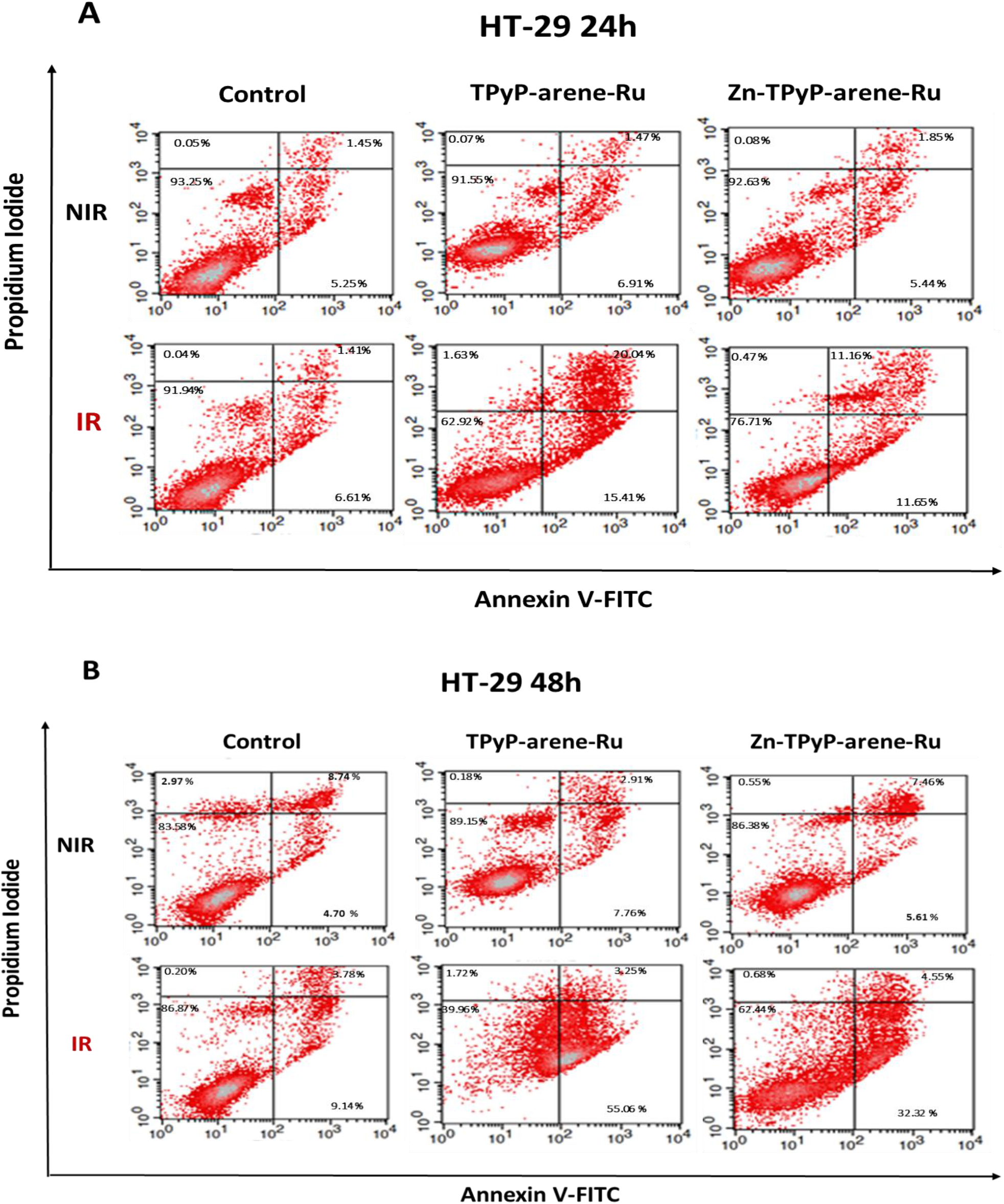
Effects of photoactivation of PS-arene-Ru on HT-29 cell line apoptosis. Cells were grown for 36h in an appropriate culture medium before exposure or not to TPyP or Zn-TPyP-arene-Ru complexes. After 24h incubation, cells were irradiated or not with a 630-660 nm CURElight lamp at 75 J/cm^2^ (PhotoCure ASA, Oslo, Norway). HT-29 cells were also stained, 24h post-PDT (A) and 48h-post-PDT (B) with Annexin V-FITC and PI, and apoptosis was analyzed by flow cytometry. The upper right quadrant represents the percentage of late apoptosis, and the lower right quadrant represents early apoptosis. Data are shown as mean ± SEM (n = 3).

### 3.4. TPyP and Zn-TPyP-arene-Ru induce DNA damage in CRC cells

To prove the role of TPyP and Zn-TPyP-arene-Ru-PDT on apoptosis, other investigations on later stages of this process had to be evaluated. Here, we were interested in the state of the poly-ADP-ribose polymerase (PARP). PARP is an enzyme engaged in DNA repair. Cleavage of PARP is considered as a hallmark of cells undergoing apoptosis. We compared the expression of native and cleaved PARP forms in treated and untreated cells using Western blotting. After PDT, results showed that TPyP and Zn-TPyP-arene-Ru induced PARP cleavage as shown by the highly apparent 89 kDa cleavage fragment for HCT116 (Figures 9B and 10B) and HT-29 (Figures 11B and 12B) cell lines, associated with a decreased expression of the native PARP in treated cells compared to control at 24 and 48h.

**Figure 9:**
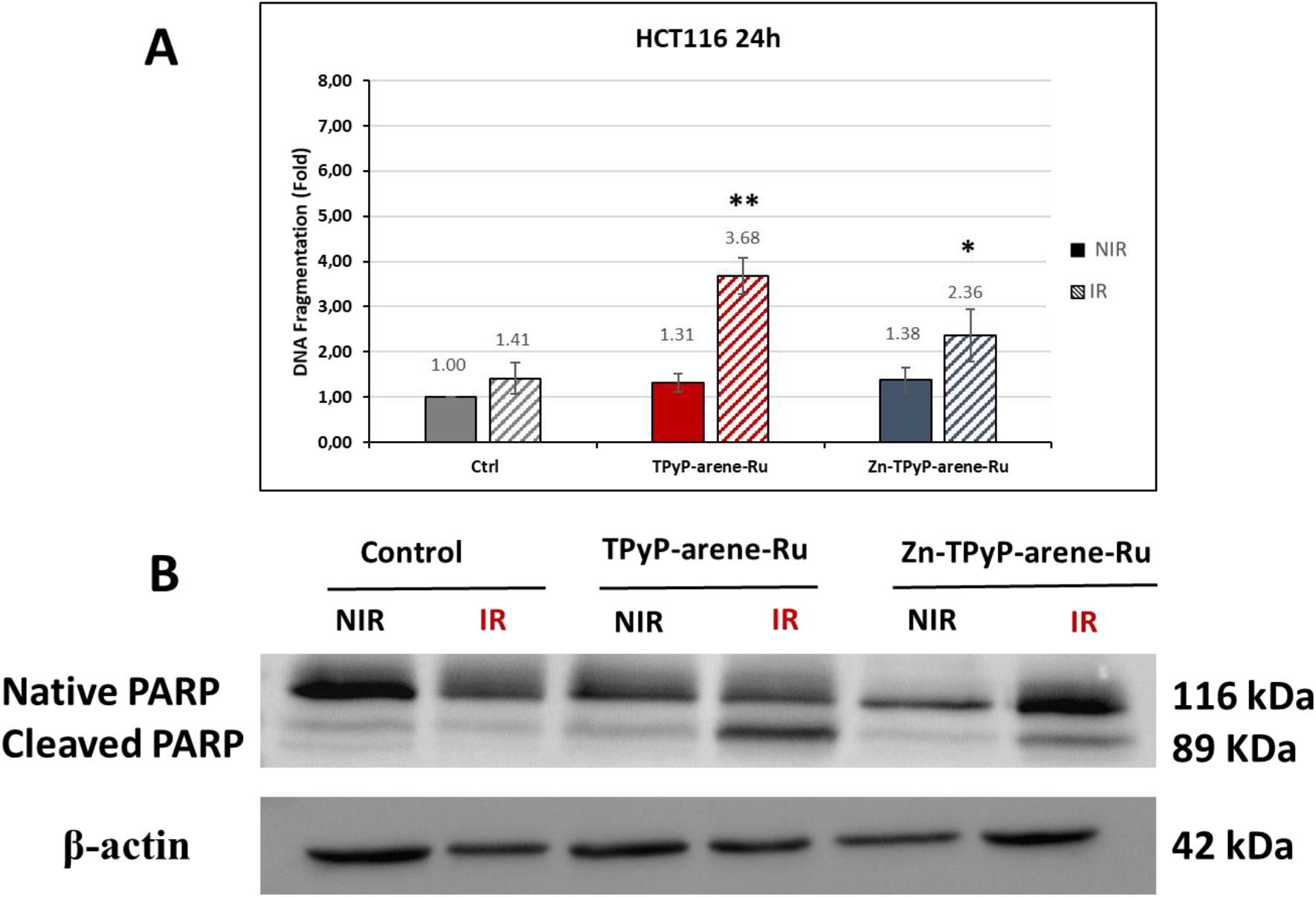
Effects of photoactivation of PS-arene-Ru on apoptosis-related protein expression and DNA fragmentation in colorectal HCT116 cells at 24h. Cells were grown for 36h in an appropriate culture medium before exposure or not to TPyP or Zn-TPyP-arene-Ru complexes. After 24h incubation, cells were irradiated or not with a 630-660 nm CURElight lamp at 75 J/cm^2^ (PhotoCure ASA, Oslo, Norway). (A) Total lysates were harvested and expression of PARP apoptosis-related protein was determined by Western blot analysis. β-actin was used as a loading control. Blots shown are representative of three separate experiments. (B) DNA fragmentation in HCT116 cells 24h post-PDT was quantified from cytosol extracts by ELISA. Results are reported as n-fold compared to control. Data are shown as mean ± SEM (n= 3). \**p* < 0.05 ** and *p* < 0.01.

**Figure 10:**
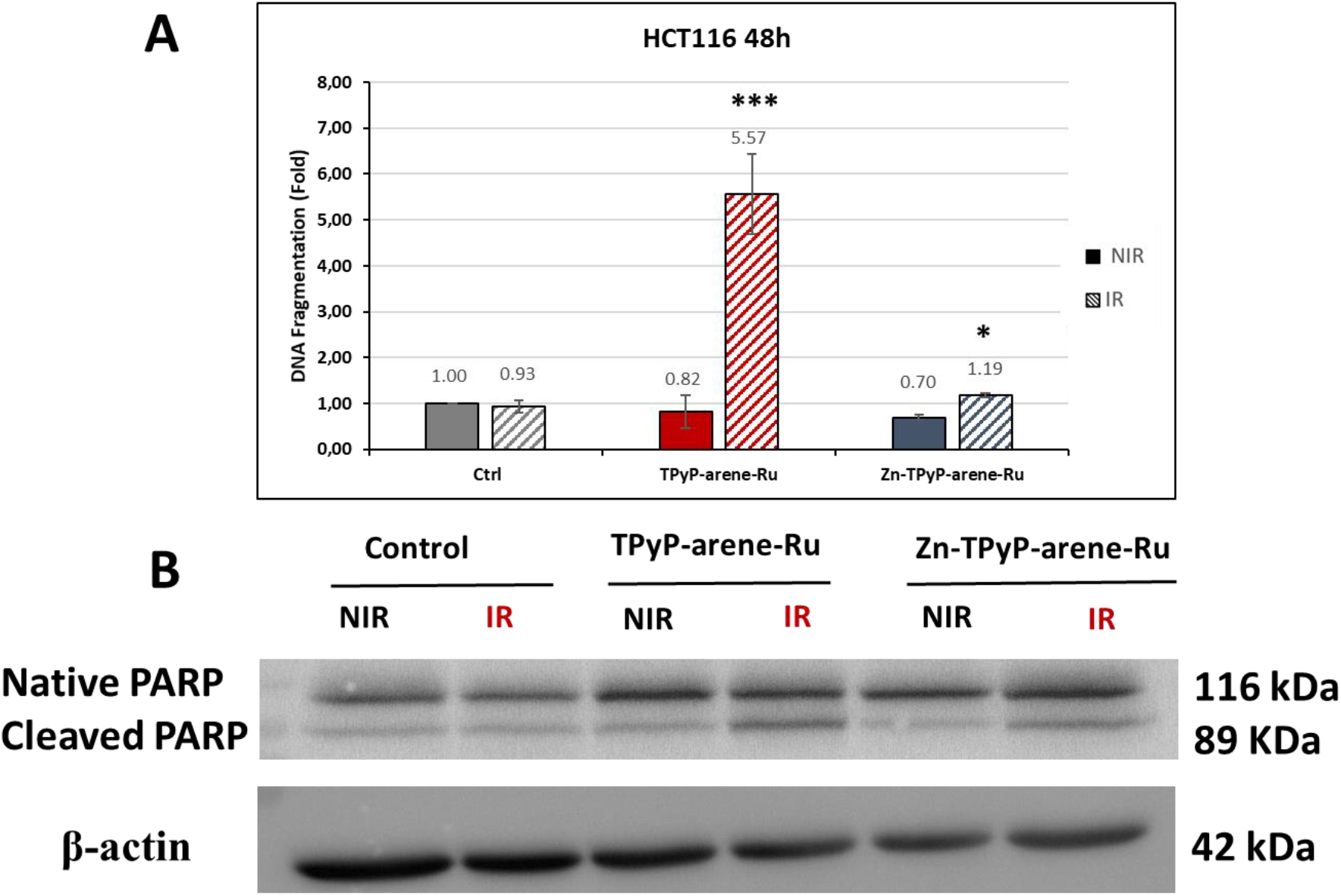
Effects of photoactivation of PS-arene-Ru on apoptosis-related protein expression and DNA fragmentation in colorectal HCT116 cells at 48h. Cells were grown for 36h in an appropriate culture medium before exposure or not to TPyP or Zn-TPyP-arene-Ru complexes. After 24h incubation, cells were irradiated or not with a 630-660 nm CURElight lamp at 75 J/cm^2^ (PhotoCure ASA, Oslo, Norway). (A) Total lysates were harvested and expression of PARP apoptosis-related protein was determined by Western blot analysis. β-actin was used as a loading control. Blots shown are representative of three separate experiments (B) DNA fragmentation in HCT116 cells 48h post-PDT was quantified from cytosol extracts by ELISA. Results are reported as n-fold compared to control Data are shown as mean ± SEM (n = 3). \**p* < 0.05 and *** *p* < 0.001.

**Figure 11:**
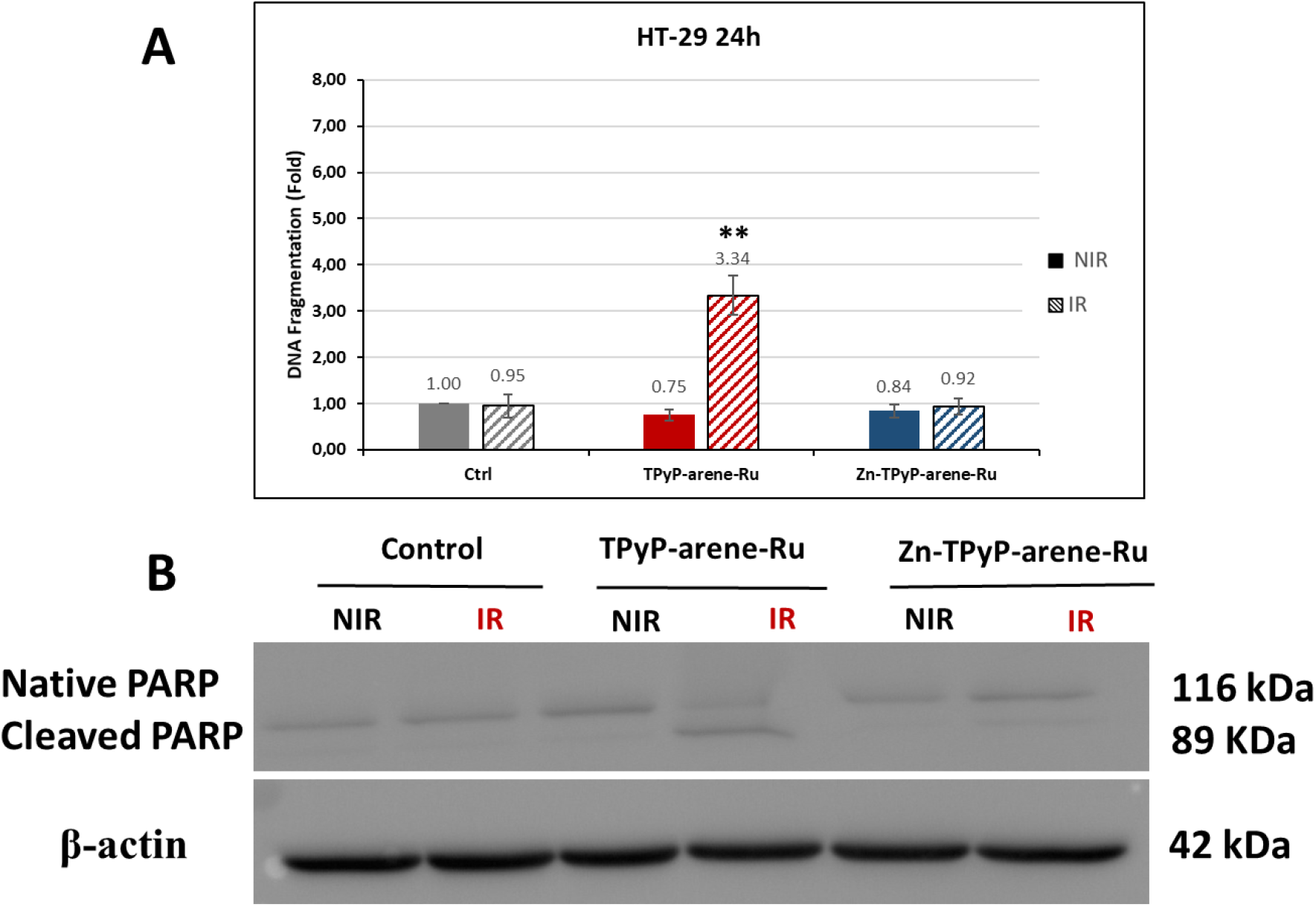
Effects of photoactivation of both compounds on apoptosis-related protein expression and DNA fragmentation in colorectal HT-29 cells at 24h. Cells were seeded in T-25 culture flasks and grown for 36 h in an appropriate culture medium before exposure or not to TPyP or Zn-TPyP-arene-Ru. After 24 h incubation, cells were irradiated or not with a 630-660 nm CURElight lamp at 75 J/cm^2^ (PhotoCure ASA, Oslo, Norway). (A) DNA fragmentation in HT-29 cells 24h post-PDT was quantified from cytosol extracts by ELISA. Results are reported as n-fold compared to control. (B) Total lysates were harvested and expression of PARP apoptosis-related protein was determined by Western blot analysis. β-actin was used as a loading control. Blots shown are representative of three separate experiments Data are shown as mean ± SEM (n = 3). ** *p* < 0.01.

**Figure 12:**
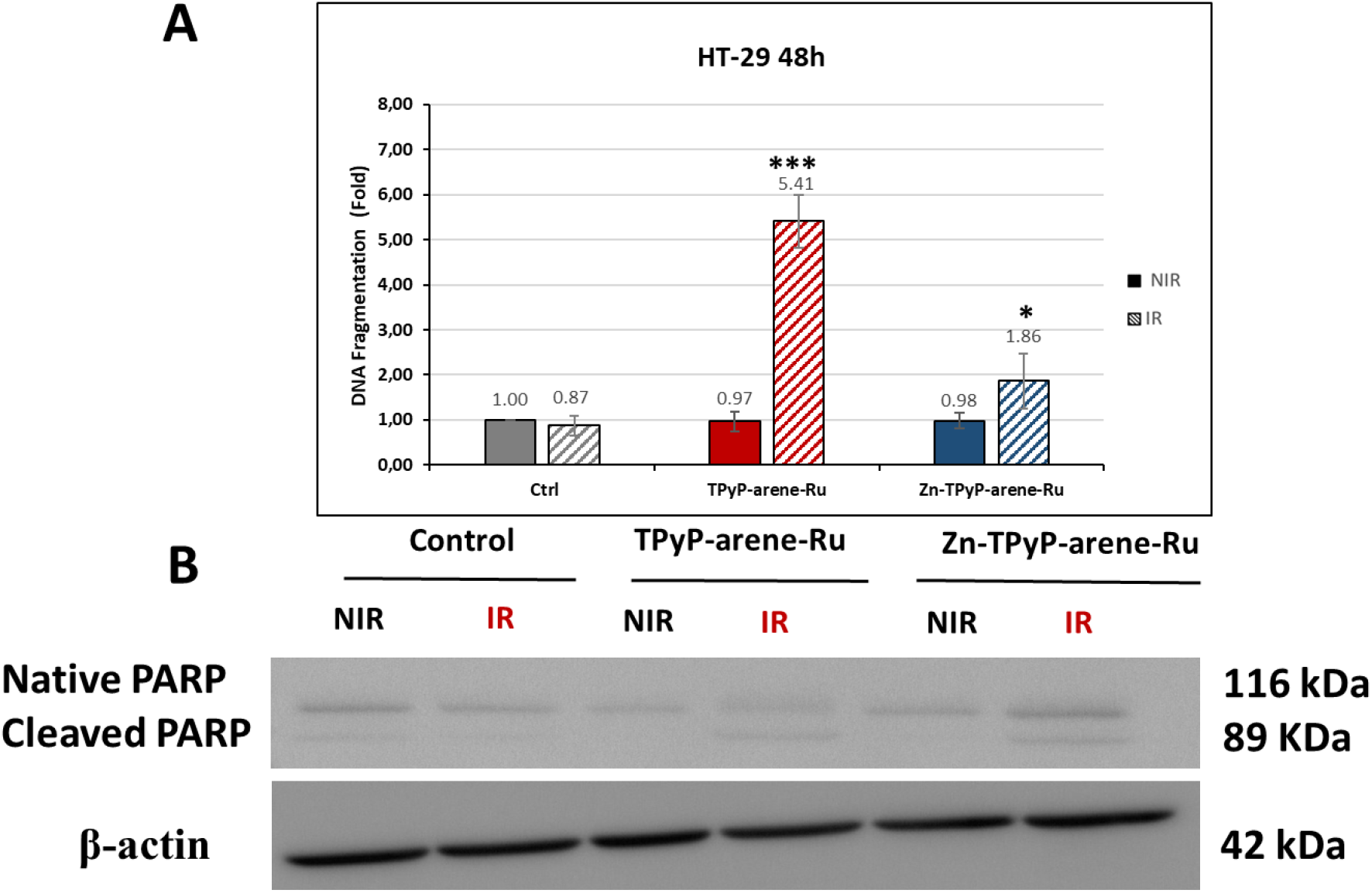
Effects of photoactivation of both compounds on apoptosis-related protein expression and DNA fragmentation in colorectal HT-29 cells at 48h. Cells were seeded in T-25 culture flasks and grown for 36 h in an appropriate culture medium before exposure or not to TPyP or Zn-TPyP-arene-Ru. After 24 h incubation, cells were irradiated or not with a 630-660 nm CURElight lamp at 75 J/cm^2^ (PhotoCure ASA, Oslo, Norway). (A) DNA fragmentation in HT-29 cells 48h post-PDT was quantified from cytosol extracts by ELISA. Results are reported as n-fold compared to control. (B) Total lysates were harvested and expression of PARP apoptosis-related protein was determined by Western blot analysis. β-actin was used as a loading control. Blots shown are representative of three separate experiments Data are shown as mean ± SEM (n =3). \**p* < 0.05 and \*\*\**p* < 0.001.

Furthermore, to study the nuclear changes in apoptosis caused by TPyP and Zn-TPyP-arene-Ru treatments with PDT, DNA fragmentation was evaluated by ELISA assay in both cell lines at 24 and 48h. The outcomes indicate that in HCT116 cells, TPyP-arene-Ru leads to a significant increase in DNA fragmentation by 3.68-fold at 24h and 5.57-fold at 48h compared to non-irradiated conditions 1.31-fold and 0.82-fold respectively compared to control. Similarly, Zn-TPyP-arene-Ru coupled with PDT increased DNA fragmentation by 2.36-fold and 1.19-fold at 24 and 48h respectively compared to non-irradiated conditions 1.38-fold and 0.70-fold compared to control (Figures 9A and 10A).

HT-29 showed similar results, TPyP-arene-Ru coupled with PDT induce a significant increase in DNA fragmentation by 3.34-fold at 24h and 5.41-fold at 48h, whereas the non-irradiated condition showed no significant effect with 1.31-fold and 0.82-fold respectively compared to control. Likewise, Zn-TPyP-arene-Ru with PDT increased DNA fragmentation mainly at 48h by 1.86-fold compared to non-irradiated condition 0.98-fold compared to control (Figures 11A and 12A).

## 4. Discussion

PDT is an innovative cancer therapy that offers advantages over conventional treatments. Nevertheless, despite its potential advantages, only a small number of PS are approved like porphyrin-types. This type of PS is useless by itself due to its low solubility in biological media, but when incorporated into metallizations, it can be internalized into cells. Based on this theory the conjugation of porphyrin with metals has received much attention [45].

Recent scientific studies showed that ruthenium complexes are one of the most interesting metal-based drugs used in the treatment of several cancers such as colon cancer [29,46]. Porphyrin and its derivatives are one of the most important Ru-conjugated PSs. The interesting properties of Ru complexes have led to their potential use in various fields such as PS and photoactive DNA cleavage agents for therapeutic purposes [47]. Recently, the attempt to find alternative approaches to enhance the specificity and selectivity of drug delivery to cancer cells has received a great deal of attention [35]. To overcome the solubility and stability problems of Ru-based PS, metal assemblies have begun to receive considerable interest, as drug encapsulation carriers [48–50].

Several studies reported that porphyrin-Ru complexes had significant anticancer effects. Bogoeva et al. reported Ru porphyrin-induced photodamage in bladder cancer cells [51]. In addition, Schmitt et al. demonstrated that five 5,10,15,20-tetra(4-pyridyl)porphyrin (TPP) arene Ru (II) derivatives and a p-cymeneosmium and two pentamethylcyclopentadienyliridium exhibited excellent phototoxicities toward melanoma cells when exposed to laser light at 652 nm [41]. Cellular uptake and localization microscopy studies of [Ru_4_-(η_6_-C_6_H_5_CH_3_)_4_(TPP)Cl_8_] and [Rh_4_-(η_5_-C_5_Me_5_)_4_(TPP)Cl_8_] revealed that they accumulated in the melanoma cell cytoplasm in granular structures different from lysosomes. Another study provided by Rani-Beeram et al. established that fluorinated Ru porphyrin presented a strong DNA interaction that leads to its cleavage in melanoma cells [52]. Most recently we reported that in cubic or prismatic cage can serve as an ideal carrier for PS to treat rheumatoid arthritis [37]. Furthermore, we demonstrated in another study that the combination of tetrapyridylporphin and arene Ru (II) complexes plays a crucial role as a PDT-conjugated anticancer agent for the treatment of synovial sarcoma [53].

In the current study, we are the first to use TPyP-arene-Ru cubic metallacages to treat CRC. We wanted to demonstrate the potential of these cages as a drug delivery system for tetrapyridylporphin PS. For this purpose, we evaluated the effect of TPyP-arene-Ru complex associated with PDT on two human CRC cell lines HCT116 and HT-29. As we explained before, our cages contain metal-ligand so to evaluate the effect of metal on our PS, a diamagnetic metal (Zn^2+^) was introduced in the center of the tetrapyrrole ring in the TPyP panels (Zn-TPyP).

First of all, we evaluated the photocytotoxicity of TPyP and Zn-TPyP-arene-Ru, and we demonstrated that both structures have a dramatic significant cytotoxic effect on the two CRC cell lines studied. This result was expected due to the distinct structure of these cages containing two units of tetrapyridylporphin as PS, which has the effect of strengthening the effectiveness of the treatment and consequently allows us to reduce the dose necessary to obtain a better therapeutic effect. TPyP-arene-Ru has shown to induce remarkable cytotoxicity by 5 to 8-folds when compared to the Zn-tetrapyridylporphin analog on both cell lines. We can relate this to the higher fluorescence of PS with a metallic center. This occurs because when the PS is irradiated it causes its excitation, that is, it absorbs part of the irradiation energy and reaches a higher energy state (excited singlet state). Fluorescence is a consequence of the energetic decay from the excited state of the PS to the minimum energy state. Therefore, high fluorescence quantum yield (ØF) suggests that much of the energy in the singlet excited state of the PS returns to the ground state without passing through the triplet excited state. Consequently, generating more fluorescence but leaving behind less energy in the triplet state to interact with O_2_ and give rise to ROS leading to cell death which decreases the treatment efficacity. However, in all cases, we systematically obtained a significant cytotoxic effect for which the determined IC_50_ values are very low, in the nanomolar range. Secondly, another interesting result, we demonstrated that tested metallacages have no cytotoxic effect in the dark even with very high concentrations. These results can confirm that these two complexes can be ideal PS for PDT and these cages can give rise to significant differences in PDT effects.

Inhibition of cancer cell proliferation by cytotoxic drugs could be the result of induction of apoptosis or cell cycle arrest or a combination of both processes [54]. We investigated the possible cell growth mechanism inhibition by flow cytometry analysis. We demonstrated that TPyP and Zn-TPyP-arene-Ru coupled with PDT led to marked increases in the proportion of apoptotic cells, as reflected by the sub-G1 peaks. Whereas, both complexes have no significant effect on cell cycle phases, which can affirm that cell growth inhibition does not depend on cell cycle arrest [55]. In order, to evaluate the induction of apoptosis mechanism leading to cell death, we investigated the percentage of phosphatidylserines externalization of apoptotic cells by annexin-V-FITC/PI dual staining assay by flow cytometry. We established that photoactivation of TPyP and Zn-TPyP-arene-Ru increased dramatically the cumulative rate of early and late apoptosis which can confirm cell death via apoptosis. These results are in agreement with a study held by Silva et al. reporting the apoptotic cell death in human colon carcinoma HCT116 cells treated with Ru(II)-thymine complex [56]. Furthermore, to validate the apoptotic mechanism, we evaluated the last stage in the death mechanism related to DNA fragmentation, we demonstrated that TPyP and Zn-TPyP-arene-Ru coupled with PDT induced significant PARP cleavage and DNA fragmentation. This result can be related to the strong potential of Ru complexes’ photophysical, photochemical properties that allow them to bind to DNA and induces its cleavage by photoactivation [57]. These demonstrated results agree with Lu et al. study which reports the anticancer effect of Ru complexes on hepatocellular carcinoma [58].

## 5. Conclusion

In this study, we evaluated for the first time the anticancer efficacy of the TPyP and Zn-TPyP arene-Ru metal assemblies on two human CRC cell lines HCT116 and HT-29. Following our hypothesis concerning the interest in porphyrin delivery, we demonstrated the strong *in vitro* anticancer efficacy of TPyP and Zn-TPyP-arene-Ru complexes coupled with PDT. Our results showed significant phototoxicity of both metallacages with a more remarkable effect for the devoid of the metal in its tetrapyrrole ring center. The results confirm cell death via the apoptotic pathway. These metal assemblies have proven to be ideal delivery systems for porphyrin-based PS giving a great advantage as a new therapeutic agent for anti-cancer PDT.

## Author Contributions

Conceptualization, J.M, B.L.; Methodology, J.M., A.P., L.P.; Validation, B.L., Writing original draft preparation, J.M. All authors have read and agreed to the published version of the manuscript.

## Acknowledgments

Authors are grateful to the BISCEm platform of ΩHealth Institute at Limoges University especially Catherine OUK with assistance for flow cytometry analysis.

## Abbreviation list

ATCC: American Type Culture Collection
CO_2_: Carbon dioxide
CRC: colorectal cancer
DMEM: Gibco Dulbecco’s Modified Eagle Medium
DMSO: Dimethyl sulfoxide
DNA: deoxyribonucleic acid
FBS: Fetal Bovine Serum
ØF: Fluorescence Quantum Yield
HEPES: 4-(2-hydroxyethyl)-1-piperazine ethane sulfonic acid
H_2_O_2_: Hydrogen peroxide
HRP: Horseradish Peroxidase
IARC: International Agency for Research on Cancer
IC_50_: median inhibiting concentration
kDa: kilodalton
MTT: 3-(4,5-dimethylthiazol-2-yl)-2,5-diphenyltetrazolium bromide
NaCl: sodium chloride
^1^O_2_: singlet oxygen
PARP: Poly-ADP-ribose polymerase
PBS: Phosphate-buffered saline
PDT: Photodynamic therapy
PI: Propidium Iodide
PS: photosensitizer
PVDF membrane: polyvinylidene fluoride membrane
RIPA: Radio immuno precipitation Assay
ROS: reactive oxygen species
RPMI: Roswell Park Memorial Institute medium
Ru: Ruthenium
SDS-PAGE: electrophoresis in polyacrylamide gel containing sodium dodecyl sulfate
SEM: Standard Error of the Mean
TPyP: Tetrapyridylporphin
WHO: World Health Organization
Zn: Zinc.

